## Supplementary material for "Single caudate neurons encode temporally discounted value for formulating motivation for action": Figure supplement

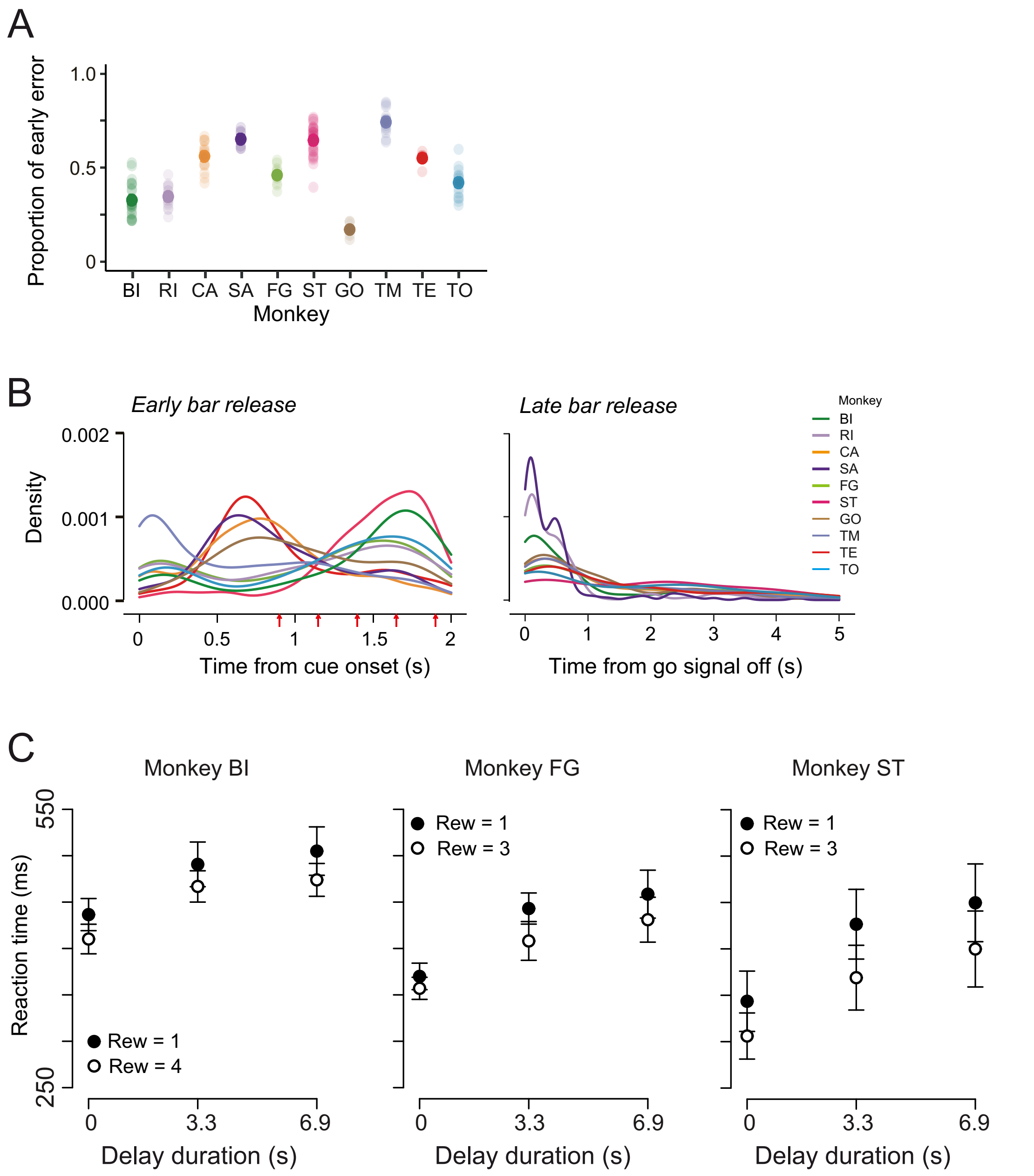


**Figure 1—figure supplement 1.** Error type and timing, and reaction time and eye position. (A) Proportion of early error for each monkey. Thick and thin dots indicate mean and data of each session, respectively. (B) Distribution of timing for early and late bar release for each monkey. Red arrows indicate the timing of go. (C) Reaction time (mean ± SD) of delayed reward task as a function of delay duration in monkeys BI, FG, and ST. Black and white symbols indicate small (1 drop) and large reward (3 or 4 drops), respectively.

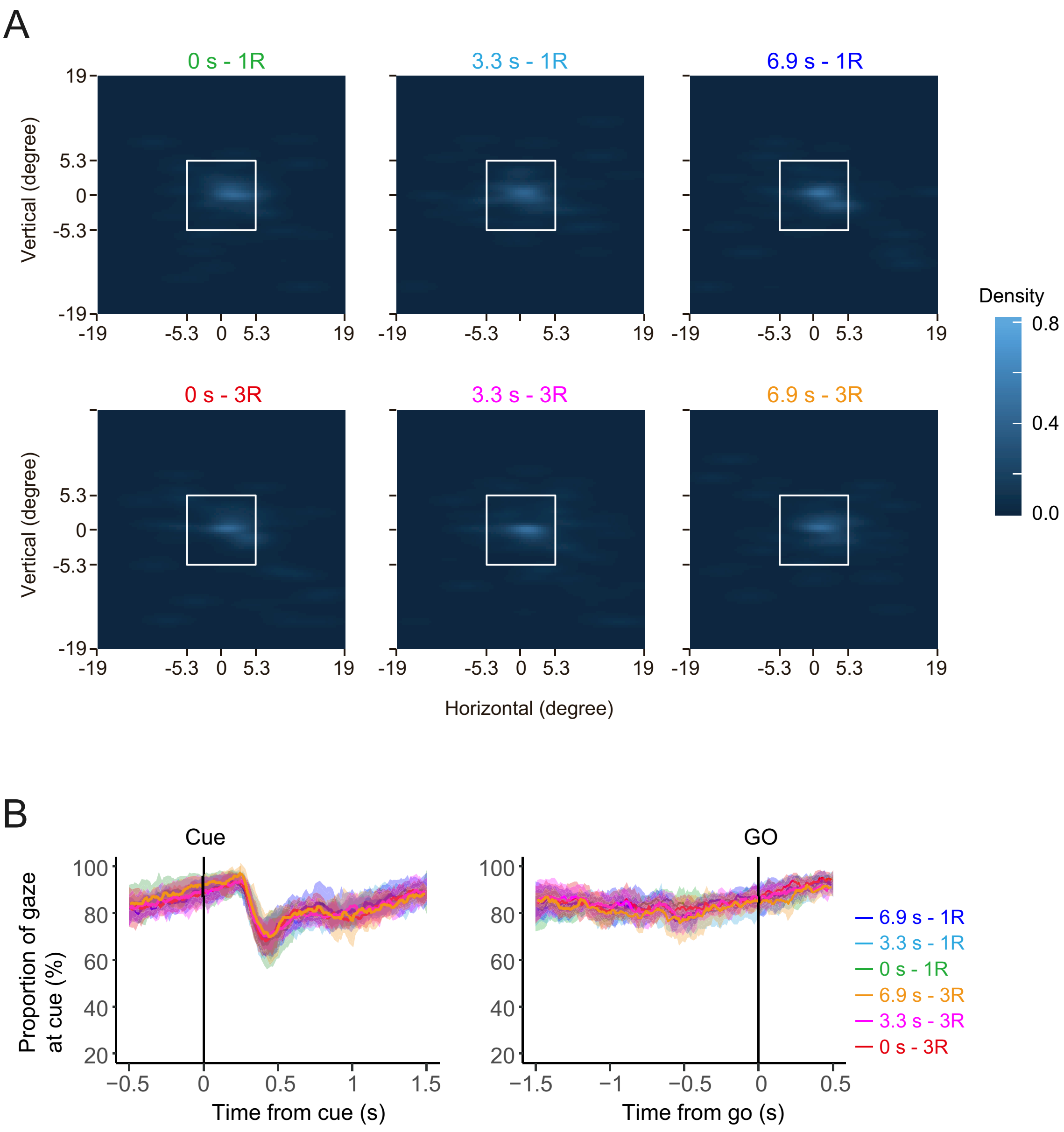


**Figure 1—figure supplement 2.** Eye position during cue period.

(A) Density plots of eye position during cue period of delayed reward task obtained from monkey RI. Colors indicate normalized looking-time. White squares indicate the frame of cue stimulus. (B) Time course of the proportion of eye position within the cue area aligned by CUE (left) and GO onset (right). Thick curves and shaded areas represent mean and SD, respectively. Colors represent rewarding condition.

| Model | Formula | Neurons |
|---|---|---|
| Model1: | $fr \sim dv$ | 5 |
| Model2: mixed effect on both slope and intercept | $fr \sim dv + (dv\|trial)$ | 1 |
| Model3: mixed effect on intercept | $fr \sim dv + (1\|traial)$ | 5 |
| Model4: mixed effect on slope | $fr \sim dv + (0 + dv\|traial)$ | 11 |
| Total | | 22 |

**Figure 3—figure supplement 1.** Error trial analysis.

Table shows that the number of neurons whose activity is explained best by models 1–4. Note that linear mixed model (LMM) analysis was applied to 22 of 27 DV-coding neurons recorded in a session in which the monkeys made at least three error trials. fr, firing rate; dv, discounted value; trial, trial type (correct or error). Seventeen neurons were differently modulated by DV depending on whether the monkey perform correct or not, while remaining 5 were similarly modulated regardless of performance.

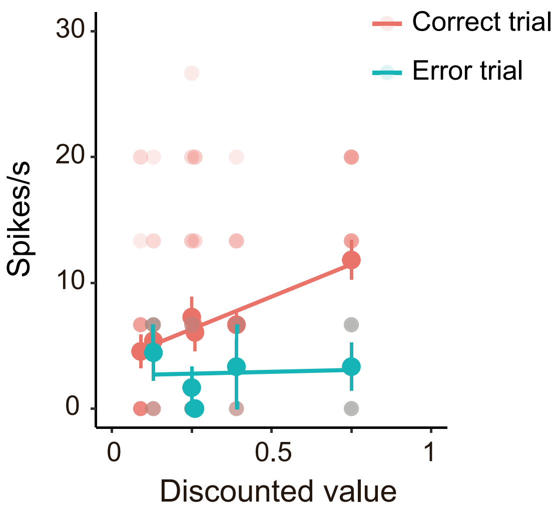


**Figure 3—figure supplement 2.** Error trial analysis.

Example of differential activity between error and correct trials of a DV-coding neuron. Thin and thick dots indicate relationship between firing rate and temporally discounted value (Equation 1) in individual trials and mean values for each rewarding condition, respectively. Colors indicate correct (red) and error (green) trials, respectively. Thick lines indicate best-fit of LMM (model 4 in Figure 3–figure supplement 1).

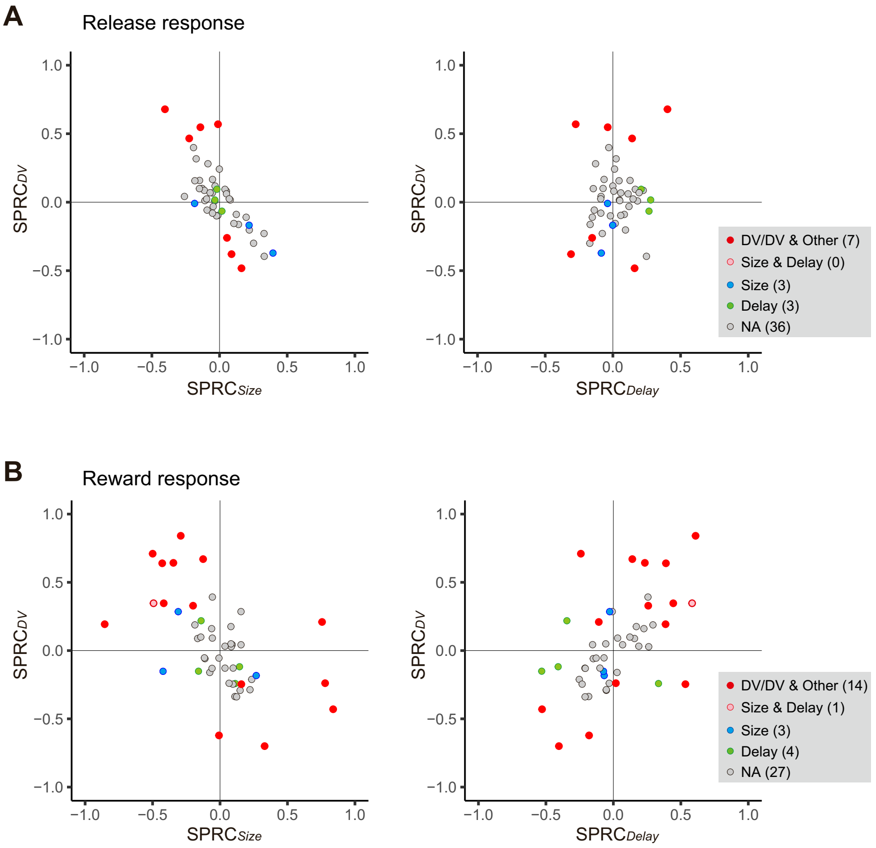


**Figure 4—figure supplement 1.** Impact of DV and comparison with delay and size on release and reward response.

(A) Scatterplot of standardized partial regression coefficients (SPRC) of DV (ordinate) against those of size and delay on release response, respectively (abscissa). (B) Same as A, but for reward response. Colored dots indicate neurons with significant ($p < 0.05$) coefficient, while gray dots correspond to neurons without any significant effect (NA). DV/DV & Other, neurons with significant coefficient of DV; Size & Delay, those with both size and delay; Size, those with size exclusively; Delay, those with delay exclusively. Numbers in parentheses indicate number of neurons.

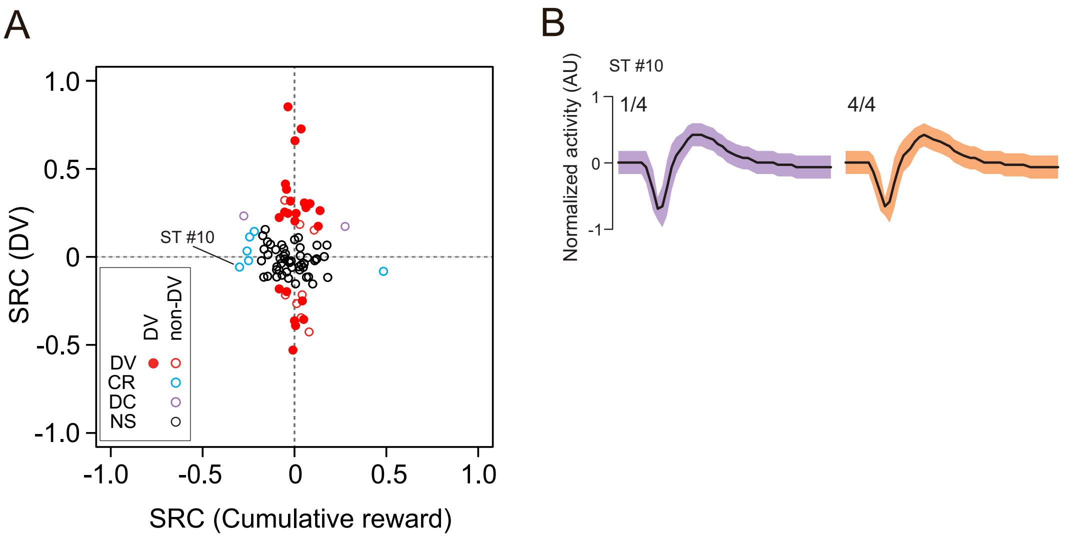


**Figure 6—figure supplement 1.**

Impact of discounted value and satiation on cue response. (A) Scatterplot of standardized regression coefficients (SRC) of discharge rates during cue period for DV (ordinate) against those for cumulative reward (abscissa). Red dots indicate DV neurons. Red and blue, and purple circles indicate non-DV neurons with significant ($p < 0.05$) coefficient for DV and cumulative reward (CR), and both respectively. Black circles correspond to neurons without any significant effect (NS). (B) Representative waveforms (mean ± SD) recorded from a CD neuron (Monkey ST #10) during first (purple) and last quartile (orange) of recording period. Changes in firing rate were not attributable to alteration in action potential isolation.

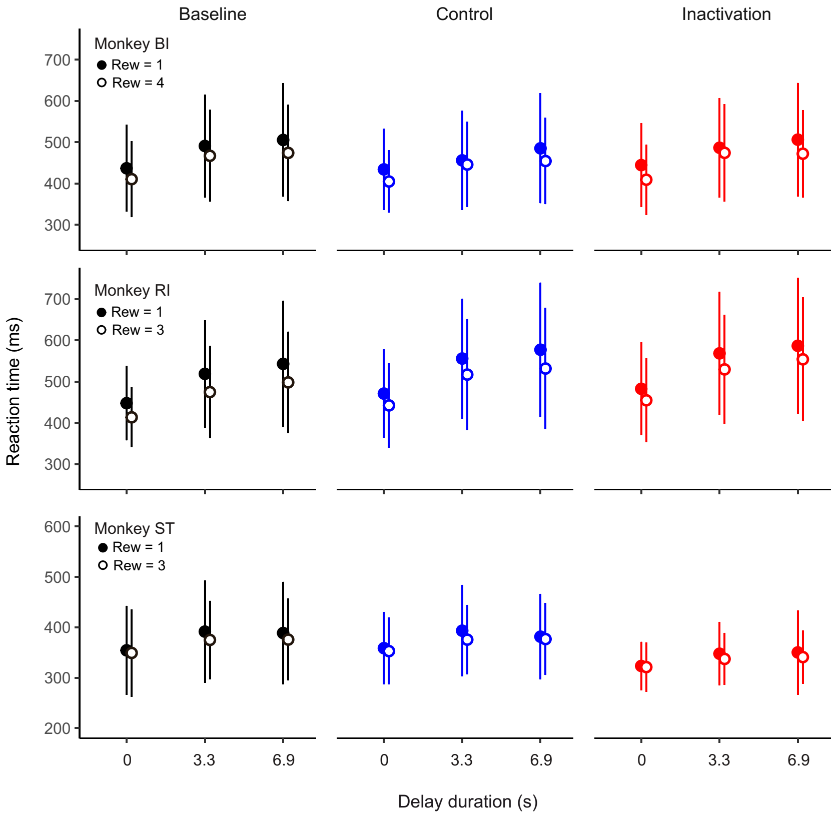


**Figure 7—figure supplement 1.** No significant effects of dCDh inactivation on reaction time in delayed reward task.

Comparison of reaction time (mean ± SD) between baseline, control and inactivation session in monkeys BI, RI and ST.

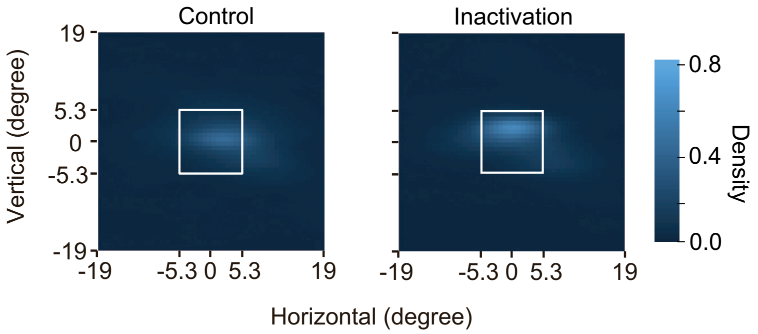


**Figure 7—figure supplement 2.** No effect of dCDh inactivation on eye position.

Density plots of eye position during cue period of delayed reward task obtained from monkey RI. Colors indicate normalized looking-time. Left and right panels for control and inactivation sessions, respectively. White squares indicate frame of cue stimulus.

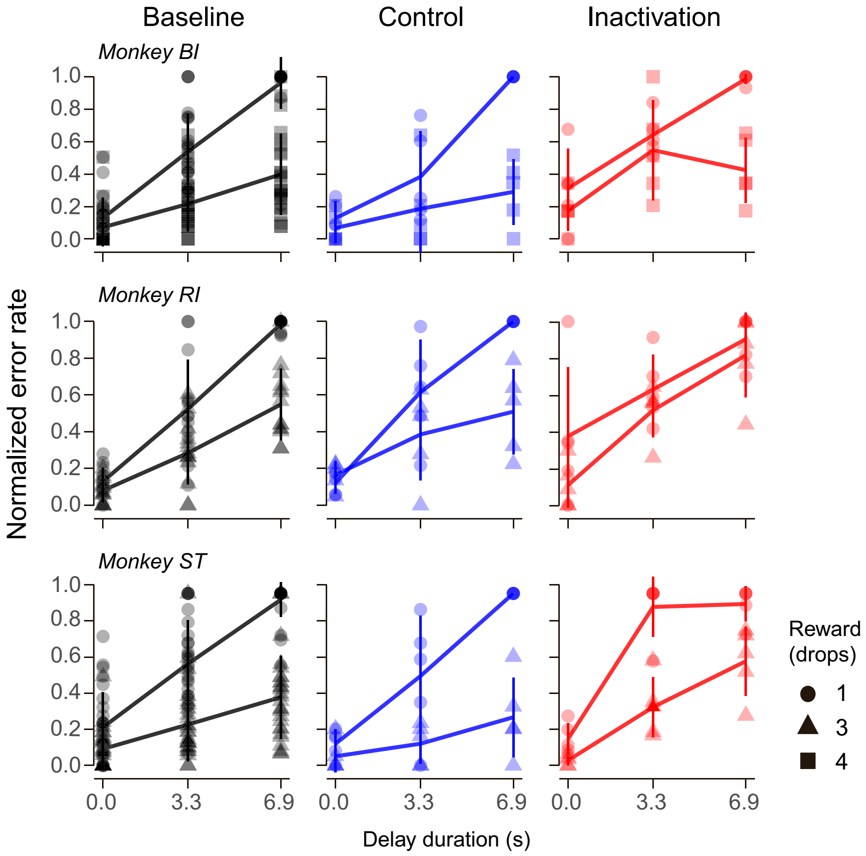


**Figure 7—figure supplement 3.** Normalized error rates in baseline, control and inactivation session of delayed reward task.

Symbols represent normalized error rates for each reward condition by maximum error rates in each session. Thick lines connect average error rates for 3 delay conditions in each reward size. Vertical lines indicate sem.

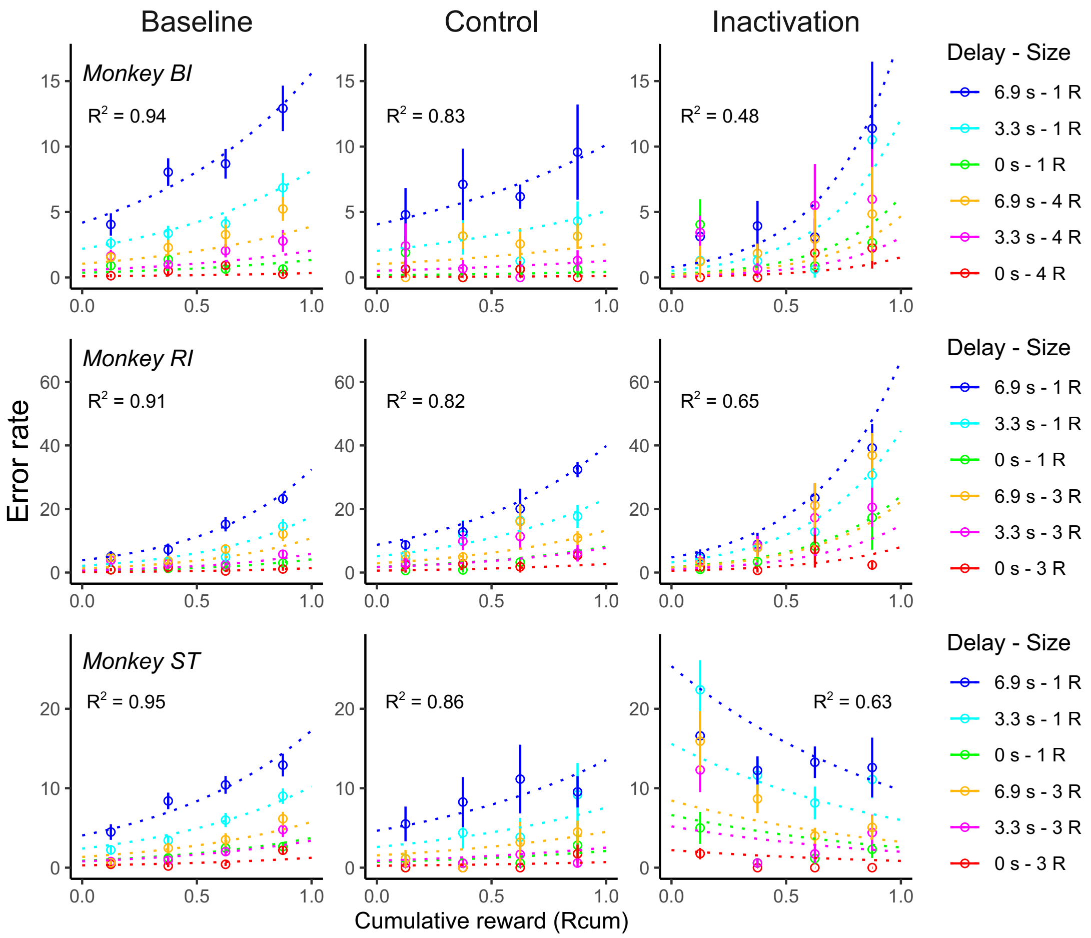


**Figure 7—figure supplement 4.** Effect of dCDh inactivation on satiation.

Error rates (mean ± sem) as a function of normalized cumulative reward ($R_{cum}$) in baseline, control and inactivation session of delayed reward task. Dotted curves are the best fit of Equation 4 to the data. Note that the satiation effect was disrupted in the inactivation session in the monkey ST, but remained normal after inactivation in the reward-size task in the same monkey (see Fig. 8C).

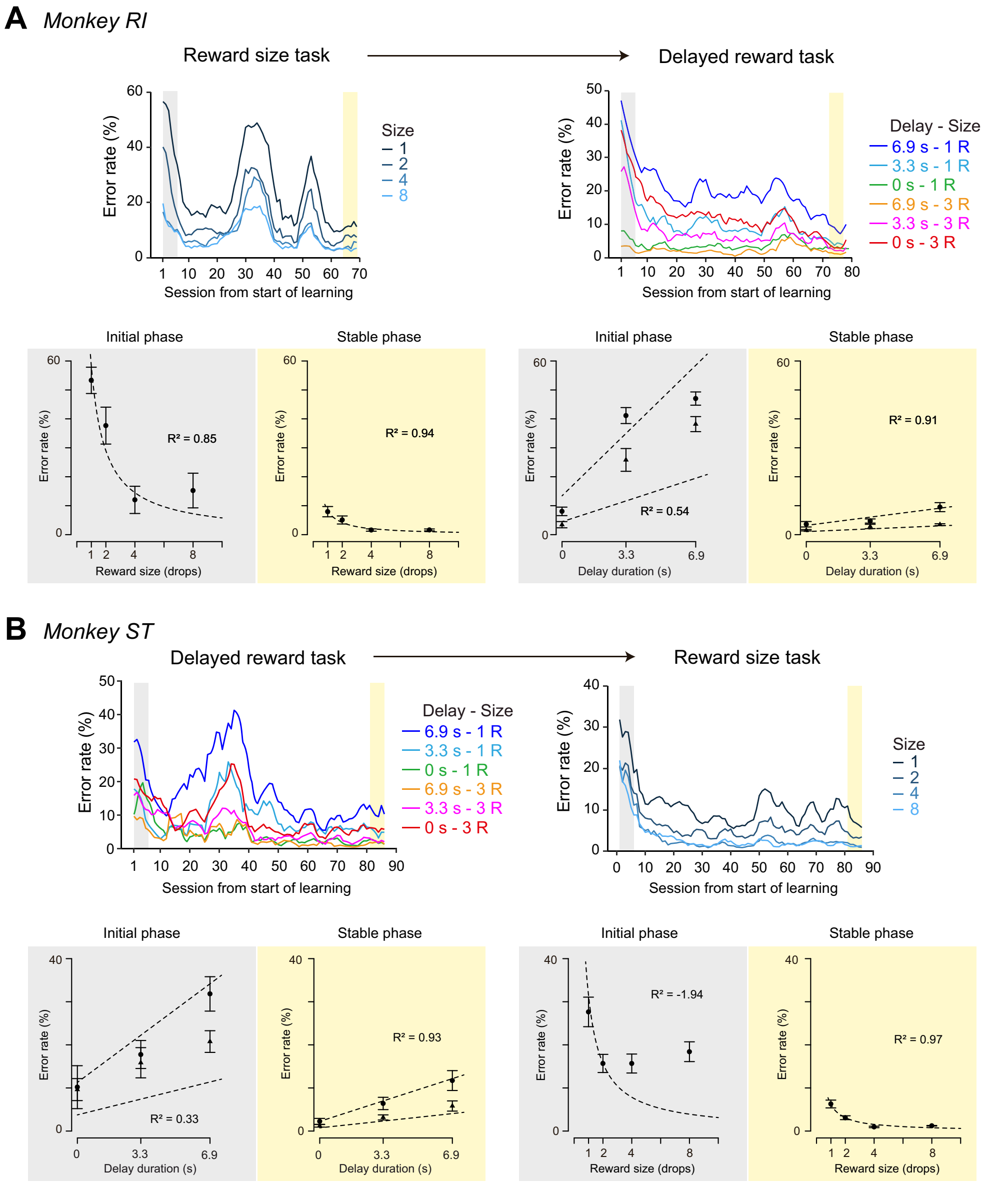


**Figure 8—figure supplement 1.** Comparison of learning in reward size and delayed reward task. (A) Monkey RI was trained with reward-size task followed by delayed reward task. (top) Error rates as a function of session were plotted for both tasks. (bottom) Error rates as a function of reward size or delay duration during initial and stable phase were shown. (B) Monkey ST was trained with delayed reward task followed by reward-size task. Others are the same as A.
